# Transposon sequencing reveals the essential gene set and genes enabling gut symbiosis in the insect symbiont *Caballeronia insecticola*

**DOI:** 10.1101/2023.09.07.556630

**Authors:** Romain Jouan, Gaëlle Lextrait, Joy Lachat, Aya Yokota, Raynald Cossard, Delphine Naquin, Tatiana Timchenko, Yoshitomo Kikuchi, Tsubasa Ohbayashi, Peter Mergaert

## Abstract

*Caballeronia insecticola* is a bacterium belonging to the *Burkholderia* genus *sensu lato*, able to colonize multiple environments like soils and the gut of the bean bug *Riptortus pedestris*. To identify the essential genome of a bacterium is a first step in the understanding of its lifestyles. We constructed a saturated *Himar1* mariner transposon library and revealed by transposon-sequencing (Tn-seq) that 498 protein-coding genes constitute the essential genome of *C. insecticola* for growth in free-living conditions. By comparing essential gene sets of *C. insecticola* and seven related *Burkholderia s.l.* strains, only 120 common genes were identified indicating that a large part of the essential genome is strain-specific. In order to reproduce specific nutritional conditions that are present in the gut of *R. pedestris*, we grew the mutant library in minimal media supplemented with candidate gut nutrients and identified several condition-dependent fitness-defect genes by Tn-seq. To validate the robustness of the approach, insertion mutants in six fitness genes were constructed and their growth-deficiency in media supplemented with the corresponding nutrient was confirmed. The mutants were further tested for their efficiency in *R. pedestris* gut colonization, confirming that gluconeogenic carbon sources, taurine and inositol, are nutrients consumed by the symbiont in the gut.

## Introduction

Essential genes in a bacterial genome are genes that are indispensable to support cellular life. Together, they constitute a minimal gene set required for a living bacterium. Essential genes are a subset of fitness genes that contribute to the reproduction of an organism independently of its environment. Thus, a fitness gene can be defined as any gene whose perturbation causes a proliferation defect and an essential gene is the extreme case when there is no proliferation at all [1, 2]. In contrast to the environment-independent essential and fitness genes, conditionally-essential or fitness genes are only required for proliferation under specific conditions. Identification of (conditionally-)essential and fitness genes of a bacterium is a first step in the understanding of its lifestyle and, from an applied point of view, it can help in designing new ligands for disease management or development of improved bio-stimulants [3, 4].

Multiple methods have been developed to identify essential and fitness genes at the whole genome scale. Among them is the systematic deletion of all possible genes and their one-by-one precise phenotyping [5, 6]. This method is laborious, making it difficult to undertake for most bacterial species. However, the recent development of Transposon-sequencing (Tn-seq) and related methods, has provided an efficient and fast solution to determine the essential genome that is accessible for a large number of bacterial species. Tn-seq allows to perform high-throughput genetic screens and to quantify in a single selection experiment the fitness impact of all genes of a genome in a condition of interest. Tn-seq uses saturated transposon-mutant populations, determines the relative abundance of all mutants in the population by high-throughput sequencing and compares these abundances in different growth conditions [7]. The analysis of the genome-wide mutation frequency in the population grown in standard conditions identifies the essential and fitness genes while the determination of altered mutation frequencies between conditions identifies the conditionally essential genes. By quantifying gene fitness by Tn-seq in many different specific environments, a fitness landscape of a bacterium in a large diversity of conditions can be created [8–10].

*Burkholderia senso lato* (*s.l.*), corresponding to the former *Burkholderia* genus and belonging to the *Betaproteobacteria*, is a large group of pathogenic, phytopathogenic, symbiotic and environmental bacteria composed of eight different genera: *Paraburkholderia*, *Robbsia*, *Pararobbsia*, *Mycetohabitans*, *Caballeronia*, *Trinickia*, *Burkholderia sensu stricto* and *Burkholderia cepacia* complex [11]. *Caballeronia insecticola* (formerly *Burkholderia insecticola*) is able to colonize multiple environments like soil or the gut of the bean bug *Riptortus pedestris* and related stinkbug species [12–14]. In this latter environment, *C. insecticola* is involved in a mutualistic relationship with the insects. The symbiotic bacteria colonize an exclusive niche composed of crypts located in the posterior midgut region of the insect where, starting from a limited number of infecting bacteria acquired from the environment through feeding, a massive extracellular population is established in the midgut crypts in a few days [15, 16]. In return, *C. insecticola* enhances the insect’s development, reproduction and immunity [17, 18]. A transcriptome analysis of *C. insecticola* in free-living growth and during colonization of the *R. pedestris* gut crypts suggested that *C. insecticola* is fed in the gut with specific nutrients and also recycles host metabolic wastes, and in return, the bacterial symbiont provides the host with essential nutrients limited in the insect food, contributing to the rapid growth and enhanced reproduction and immunity of the bean bug host [13].

Here, a saturated *Himar1* mariner transposon mutant library of *C. insecticola* allowed us to identify the essential gene set for *in vitro* growth in standard rich medium and in a defined minimal medium. In addition, to mimic specific nutrient conditions that are likely present in the insect symbiotic organ, this mutant library was grown in minimal media supplemented with specific gut nutrients and we identified several condition-dependent fitness genes. Thus identified candidate genes for growth on different carbon sources were selected for mutagenesis and the constructed mutants were tested for growth on the relevant nutrient sources and for their capacity to colonize the *R. pedestris* symbiotic gut organ in mono inoculation or in co-inoculation with the wild-type bacteria.

## Materials and methods

### Bacterial strains and growth conditions

*C. insecticola* strains were cultured at 28°C in Yeast-Glucose medium (YG: 5 g/L yeast extract; 1 g/L NaCl; 4 g/L glucose) for routine use or in minimal media (MM) supplemented with various carbon, sulphur or nitrogen sources for the Tn-seq screens (Supplementary Dataset s1). *C. insecticola* RPE75 is a spontaneous rifampicin-resistant derivative of the wild-type strain RPE64 [12]. For standard molecular microbiology purposes, *E. coli* strains DH5α, HB101, WM3064, MFDpir, S17-1λpir and their derivatives were grown in LB medium (5 g/L yeast extract, 10 g/L tryptone, 5 g/L NaCl) at 37°C. Growth of the MFDpir and WM3064 strains that are Δ*dapA* derivatives, auxotroph for diaminopimelic acid (DAP) synthesis, required the supplement of 300 µg/mL DAP to the medium. When appropriate, antibiotics were added to the medium in the following concentrations: 50 µg/mL kanamycin (Km) for *E. coli* and 30 µg/mL for *C. insecticola*; 25 µg/mL chloramphenicol (Cm); 30 µg/mL rifampicin (Rif); 100 µg/mL ampicillin (Amp). For solid agar plates, the media were supplemented with 1.5 % agar.

### Generation of a *C. insecticola* RPE75 *Himar1* transposon library

A Himar1 transposon library was generated following methods described before [19]. Detailed procedures for the construction and quality control of the library are provided in the Supplementary Text.

### Tn-seq screening of the *Himar1* transposon library for growth with different nutrients

An aliquot of the Tn-seq library was 100-fold diluted to reach a suspension of 2×10^8^ cfu/mL. Hundred µL of this dilution was inoculated into 20 mL of a growth medium, supplemented with Rif and Km, to obtain an initial inoculum of 10^6^ cfu/mL (OD_600nm_≈0.0015). Eleven growth conditions were tested: YG medium and ten different minimal media (MM) supplemented with various carbon, sulphur or nitrogen sources. The assembly from stock solutions and the composition of the MM media are provided in Supplementary Dataset s1. Cultures were incubated at 28°C, with shaking at 180 rpm. When the cultures reached an OD_600nm_≈1, corresponding to approximately 9 to 10 generations of multiplication, bacteria were collected by centrifugation at 4 000 rpm for 20 minutes at 4°C and the pellets were stored at −20°C until DNA extraction. Each condition was performed in triplicates.

### DNA extraction and preparation of the high-throughput sequencing libraries

Genomic DNA was extracted from the bacterial pellets using the MasterPure™ Complete DNA and RNA purification kit (Epicentre) according to the manufacturer’s instructions. Preparation, concentration determination and quality control of the high-throughput sequencing libraries were done following procedures described before [19] and detailed in the Supplementary Text.

### Sequencing and sequence data treatment

Sequencing and sequence data treatment were done as described [19], using the reference genome of *C. insecticola* (accession n° NC_021287.1 (chromosome 1), NC_021294.1 (chromosome 2), NC_021288.1 (chromosome 3), NC_021289.1 (plasmid 1), NC_021295.1 (plasmid 2)). Detailed information is provided in the Supplementary Text.

### Identification of (conditionally) essential genes by Transit software

Tn-seq sequencing data was handled by TRANSIT Version 3.2.0 [20] using Hidden Markov Model analysis to determine essentiality within a single condition and Resampling analysis to compare two conditions. Detailed information is provided in the Supplementary Text.

### Homology-based comparison between *Burkholderia s.l.* species

OrthoVenn2 [21] was used to compare the essential genomes of eight selected *Burkholderia* species. We used E-value of 1e −15 and inflation value of 1.5 as analysis parameters. Data from the following studies were used to establish the lists of the essential genome for each of seven other *Burkholderia* species besides *C. insecticola*: *Burkholderia pseudomallei* strain K96243 [22], *Burkholderia cenocepacia* strain J2315 [23], *B. cenocepacia* strain H111 [24], *B. cenocepacia* strain K56-2 [25], *Burkholderia thailandensis* strain E264 [26]. *Burkholderia vietnamiensis* strain LMG10929 [27] and *Paraburkholderia kururiensis* strain M130 [27]

### Construction of *C. insecticola* RPE75 insertion mutants fluorescent protein tagged strains

For insertion mutagenesis of *C. insecticola* RPE75, internal fragments (300-600 bp) of the target gene were amplified by PCR (Table s1) and cloned into the pVO155-p*nptII*-*GFP* vector using restriction enzymes *Sal*I-*Xba*I or *Xho*I-*Xba*I and ligation or alternatively by Gibson assembly cloning. Constructs were introduced in *E. coli* DH5α by heat shock transformation and selection with Km. Candidate colonies were confirmed by colony-PCR and Sanger sequencing (Eurofins Genomics). The plasmid construct was transferred to the recipient *C. insecticola* RPE75 strain by triparental conjugation, with the *E. coli* DH5α donor strain and the *E. coli* HB101.pRK600 helper strain. Transconjugants were selected with Rif and Km. Candidate *C. insecticola* mutants were verified by colony-PCR and by checking the GFP fluorescence. A mScarlett-I-tagged strain of *C. insecticola* was created by introducing a Tn7-Scarlet transposon using triparental mating as described [28, 29] (Supplementary Text).

### Bacterial growth determination on different nutrients

To determine the growth capacity of the *C. insecticola* mutants on different carbon sources, precultures of tested strains were grown in MM medium. Overnight grown cultures were diluted to an OD_600nm_=0.3 in fresh medium and grown until they reached OD_600nm_≈1. The cells were pelleted by centrifugation, resuspended to an OD_600nm_=0.05 in fresh medium with the tested carbon source. These cell suspensions were dispatched in a 96-well plates, which were incubated in a SPECTROstar Nano plate incubator (BMG Labtech). The growth of the cultures in the wells was monitored by measuring the OD_600nm_ and data points were collected every hour for 48 h. Plates were incubated at 28°C with double orbital shaking at 200 rpm. Data and growth curves were analyzed using Microsoft Excel.

### Insect rearing and inoculation tests

Insect rearing and inoculation tests were conducted as before [30]. At three and five days post inoculation, insects, at the stage of the end of the second instar nymphs or the third instar, respectively, were dissected. The colonization rate of mono-inoculated insects was estimated by fluorescent signal detection of colonizing bacteria with GFP or mScarlett-labelled fluorescent proteins. For co-inoculation experiments, M4 region samples were homogenized in PBS solution and bacteria in suspension were counted by flow cytometry, using the fluorescent tags to determine the relative abundance of the two inoculated strains (see Supplementary Text for details).

## Results

### Construction and sequencing of a Tn-seq library of *Caballeronia insecticola*

We mutagenized *C. insecticola* with a *Himar1* mariner transposon that selectively inserts in TA sites of the genome. The five replicons of the *C. insecticola* genome, chromosomes 1 to 3 and plasmids 1 and 2, present a total DNA content of 6.96 Mbp and contain 6 349 annotated genes [31]. Among the 110 735 TA sites present in the genome, 67 276 of them are located in the annotated genes. Among the 6 349 annotated genes, 6 286 possess TA sites; hence, *Himar1*-based mutagenesis covers approximately 99.01 % of genes. The genes lacking TA sites (63 genes) are short to very short open reading frames varying in length from 25 to 199 nucleotides and encoding mostly peptides of unknown function (Table 1). On the other hand, genes with TA sites have a mean of 10.6 TA sites in their sequence. Thus, as the proportion of genes without TA sites in the genome is small (0.99 %) and the large majority of genes have a high number of TA sites, it was feasible to produce a genome-wide *C. insecticola* transposon insertion library with a good coverage using the *Himar1* mariner transposon. A large-scale mutagenesis resulted in about 2.5×10^8^ independent clones, representing an over 2 000-fold coverage of all theoretically possible mutants, with 80 % of the TA sites located within a gene carrying insertions.

**Table 1.** Relevant features of the genome of *Caballeronia insecticola*.

|  | Size (bp) | N° of TA sites | N° of genes | N° of TA sites in genes | N° of genes without TA | Gene size without TA (bp) |
| --- | --- | --- | --- | --- | --- | --- |
| **Chromosome 1** | 3 013 410 | 45 986 | 2 803 | 27 310 | 34 | 25-199 |
| **Chromosome 2** | 1 465 356 | 23 340 | 1 337 | 14 103 | 18 | 53-170 |
| **Chromosome 3** | 900 830 | 14 869 | 790 | 9 368 | 3 | 106-164 |
| **Plasmid 1** | 1 275 199 | 20 575 | 1 157 | 13 056 | 8 | 61-143 |
| **Plasmid 2** | 309 692 | 5 965 | 262 | 3 439 | 0 | - |
| **whole genome** | 6 964 487 | 110 735 | 6 349 | 67 276 | 63 | 25-199 |

The mutant library was grown in three different standard growth media: a rich medium (YG) and a defined minimal medium (MM) with glucose or succinate as carbon source, and cultures were subjected to Tn-seq. The essential genome was determined using the Hidden Markov Model (HMM) in Transit software [20] and considering only the genes that are common to the three conditions as part of the essential gene set. This analysis classified the majority of the genes (92 %) as non-essential and 498 (8 %) protein-coding genes as essential (Fig. 1) (Supplementary Dataset s2). This proportion of the *C. insecticola* genome identified as its essential genome is very similar to what is found in other bacteria [32]. These 498 genes were mostly located on the chromosome 1 (464 genes or 93 % of total). Another 22 genes (4.5 %) were on chromosome 2 and only 5, 4 and 3 genes were found on chromosome 3, plasmid 1 and plasmid 2 respectively.

**Figure 1.**
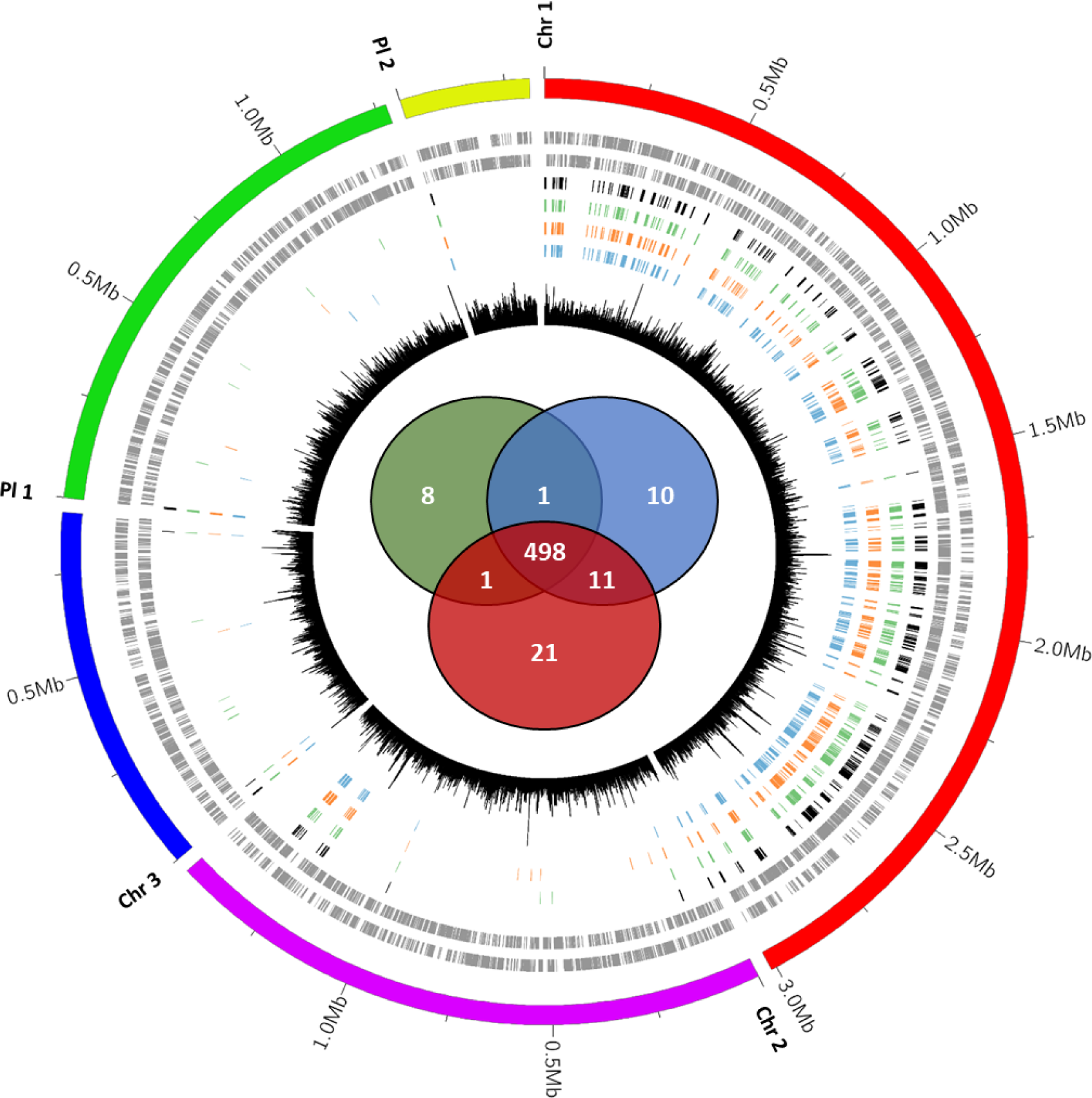
*Caballeronia insecticola* essential genes. Circular representation of the genome of *C. insecticola*, the markings outside the outer circle represent genome positions (in Mb) for each chromosome or plasmid; red is chromosome 1, purple is chromosome 2 bleu is chromosome 3, green is plasmid 1 and yellow is plasmid 2. The second and third tracks represent CDS on the forward and reverse strand, respectively. Tracks 4, 5 and 6 represent respectively essential genes (predicted by HMM Transit analysis) for YG (Green), MM glucose (Blue), MM succinate (Orange) media. Track 7 (black) shows the number of TA sites per 1 000 bp. The Venn diagram in the centre represents the number of essential genes identified for each medium: green, YG; blue, MM glucose; red, MM succinate.

According to COG (Clusters of Orthologous Genes) classification [33], the most represented category of essential genes was related to translation, ribosomal structure and biogenesis (J category) (Fig. 2). Genes encoding for 30S and 50S ribosomal proteins are examples of this functional class (Fig. 3). The cell wall biogenesis category (M category), the coenzyme transport and metabolism category (H category) and the energy conversion and production category (C category) also contain many essential functions. Examples in these three categories are genes involved in lipid A and peptidoglycan biosynthesis, the heme biosynthesis pathway or genes involved in respiration, like the ATP synthase subunits (Fig. 3) and the Respiratory Complex I subunits. Other highly represented categories are amino acid metabolism (E category) with the genes taking part in the L-histidine biosynthesis as examples or the transcription machinery (K category) with the RNA polymerase subunits. The distribution of the essential genes among the COG categories is very different compared to all the annotated genes in the genome (Fig. 2), demonstrating that our analysis identified specific functions as essential.

**Figure 2.**
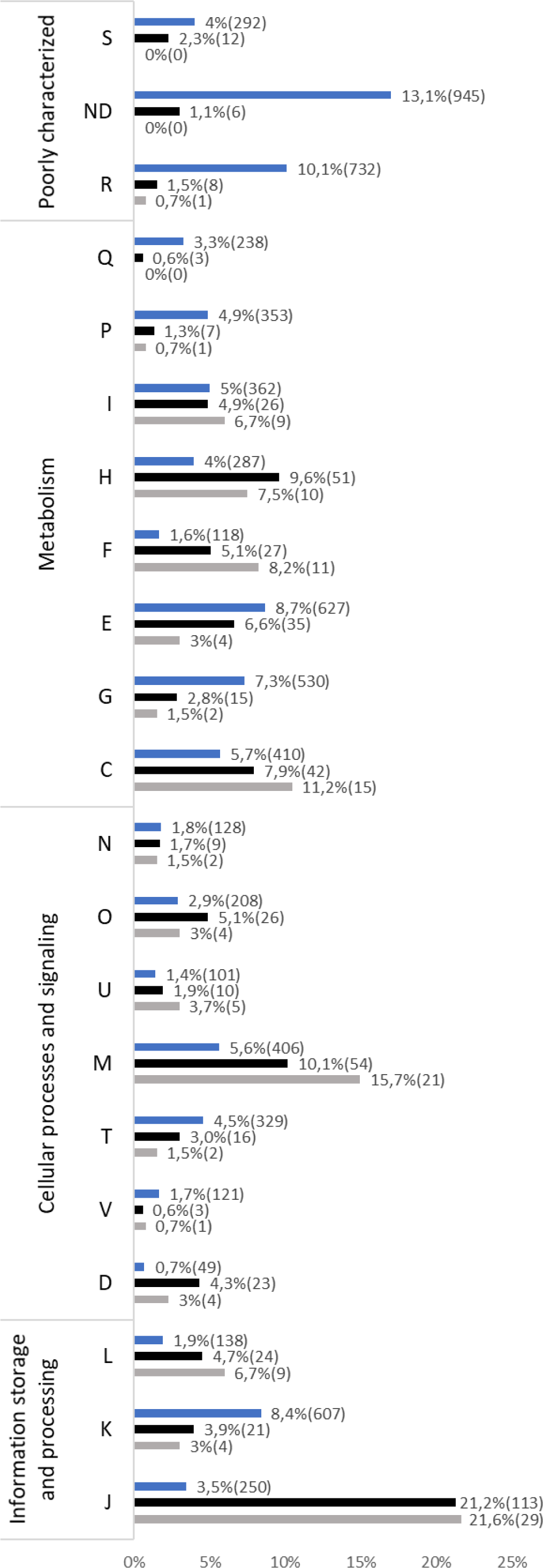
Distribution of *Caballeronia insecticola* essential genes in COG categories. The categories are S, Function unknown; ND, Not determined; R, General function prediction only; Q, Secondary metabolites biosynthesis - transport - catabolism; P, Inorganic ion transport - metabolism; I, Lipid transport - metabolism; H, Coenzyme transport - metabolism; F, Nucleotide transport - metabolism; E, Amino acid transport - metabolism; G, Carbohydrate transport - metabolism; C, Energy production - conversion; N, Cell motility; O, Post-translational modification - protein turnover – chaperones; U, Intracellular trafficking – secretion - vesicular transport; M, Cell wall - membrane - envelope biogenesis; T, Signal transduction mechanisms; V, Defence mechanisms; D, Cell cycle control - cell division - chromosome partitioning; L, Replication - recombination – repair; K, Transcription; J, Translation - ribosomal structure – biogenesis. For each category, the number of genes and the percentage that they represent are indicated. Blue histograms indicate the whole genome of *C. insecticola*, black histograms indicate the essential genome of *C. insecticola* and grey the common essential genes to eight *Burkholderia s.l.* species.

**Figure 3.**
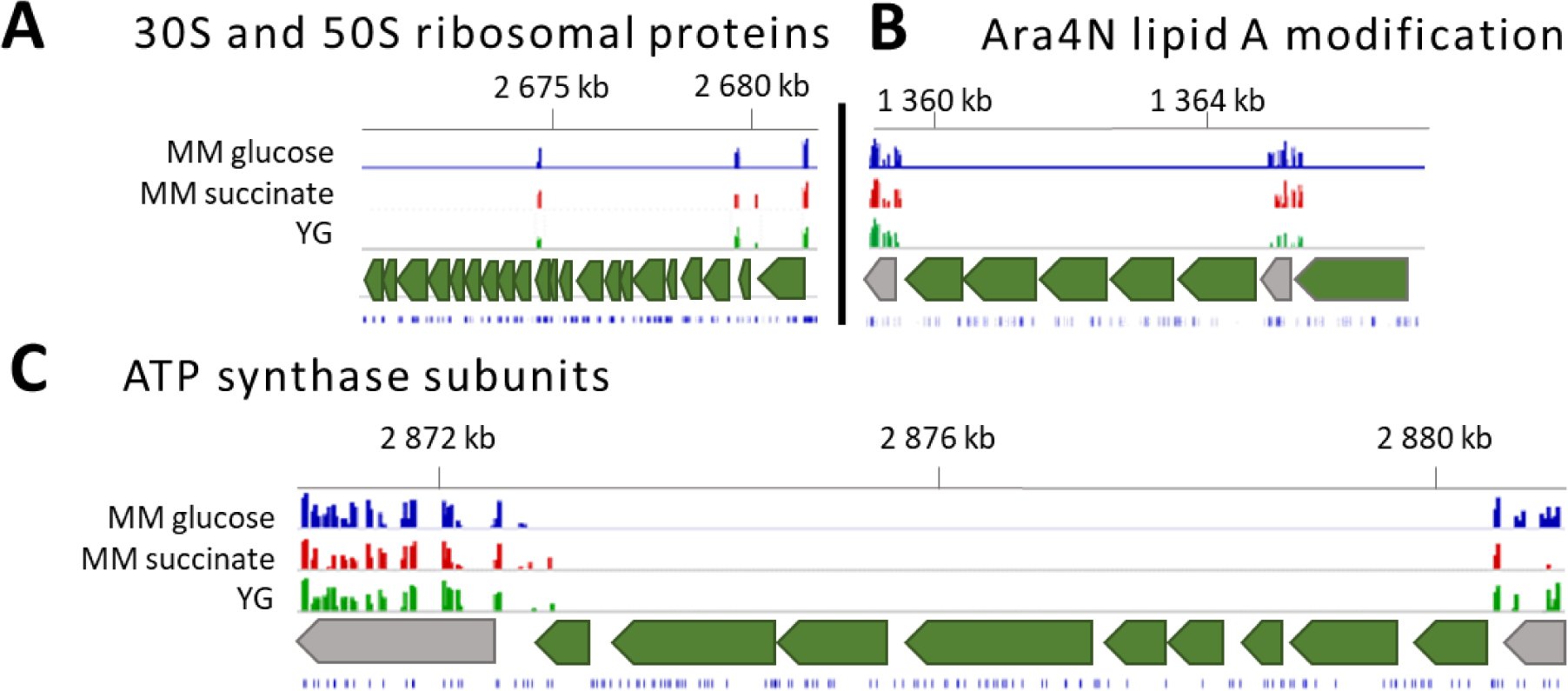
IGV plots for genomic regions containing selected *Caballeronia insecticola* essential genes. **A.** 30S and 50S ribosomal proteins encoding region. **B.** 4-amino-4-deoxy-L-arabinose (Ara4N) lipid A modification gene cluster. **C.** ATP synthase subunits encoding region. Tracks, from bottom to top: Position of TA sites, Region of interest (in green) and its flanking neighbours (in grey), Histogram of insertion counts at TA sites for the indicated experimental conditions, Genome positions (in kb) on the chromosome 1.

Of note in the cell cycle control, cell division, chromosome partitioning category (D category) are the genes involved in the replication and partitioning of the genome. Chromosome 1 has a typical organization similar to the principal chromosome of *B. cenocepacia*, carrying on the replication origin locus the genes *rpmH* (BRPE64_RS14035), *rnpA* (BRPE64_RS14030), *dnaA* (BRPE64_RS00005), *dnaN* (BRPE64_RS00010) and *gyrB* (BRPE64_RS00015), and on a nearby locus the chromosome partitioning genes *parA* (BRPE64_RS13400) and *parB* (BRPE64_RS13395) [34, 35]. Except for *rpmH* that has no TA sites, all these genes were found to be essential in agreement with their crucial role in the replication and thus persistence of Chromosome 1. On the other hand, chromosomes 2 and 3 have, similarly to plasmids 1 and 2, a plasmid-like replication origin locus, with its own distinct *parABS* system and a plasmid-like replication protein (chromosome 2: BRPE64_RS14050, BRPE64_RS14055, BRPE64_RS14060; chromosome 3: BRPE64_RS20740, BRPE64_RS20745, BRPE64_RS20750; plasmid 1: BRPE64_RS24690, BRPE64_RS24695, BRPE64_RS24700; plasmid 2: BRPE64_RS30485, BRPE64_RS30490, BRPE64_RS30495). Thus, the chromosomes 2 and 3 have plasmid-like features although they carry several essential genes. According to a new classification of bacterial replicons, the chromosomes 2 and 3 should be considered as “chromids” [36–38]. Interestingly, each of these *parABS* and replication protein-encoding genes were found to be essential, indicating that these replicons require their own cognate machinery for replication and partition. While the three chromosomes carry other essential genes, only the genes implicated in replication and partitioning are essential in the two plasmids. Thus the essentiality of these genes in this case does not mean essential for cell viability, but rather essential for the maintenance of the replicon. On the contrary, since the chromosomes contain other essential genes, these replicons and their replication/partitioning functions are essential for viability.

Taken together, the pathways of the *C. insecticola* essential genome highlight cellular functions, like transcription, translation, energy production, cell envelope biosynthesis and cell cycle, known to represent vital functions for bacteria [39, 40].

### Comparative analyses of essential genes in six *Burkholderia s.l.* species

Tn-seq techniques were used to identify essential genes in pathogenic *Burkholderia* species including *Burkholderia pseudomallei* strain K96243 [22], *Burkholderia cenocepacia* strain J2315 [23], *B. cenocepacia* strain H111 [24], *B. cenocepacia* strain K56-2 [25], in the plant-associated species *Burkholderia vietnamiensis* strain LMG10929 and *Parabukholderia kururiensis* strain M130 [27], as well as in the environmental species *Burkholderia thailandensis* strain E264 [26]. These studies revealed 505, 383, 339, 493, 620, 700 and 406 essential protein-coding genes in these bacteria, respectively. The comparison of these gene sets with the *C. insecticola* essential genes allowed us to identify a total of 120 essential genes shared between all eight species (Fig. s1; Supplementary Dataset s3). If essential genes shared by seven out of the eight species are considered, the number of common genes increases to 231 (Fig. s1) and the number of pairwise shared essential genes ranges from 195 to 412 genes with as a mean 291 genes (Fig. s2). Among the 37 clusters of orthologous genes that are essential in at least four of the considered species but not in *C. insecticola*, we found for five clusters that the *C. insecticola* genome lacked the corresponding gene. In 26 cases, the gene was present but non-essential and in six cases, the gene was duplicated in the genome.

Considering the 120 genes that are commonly essential to all eight species, an important part of them ensures functions related to the translation process (J category), the cell wall biosynthesis (M category), the coenzyme transport and metabolism (H category) or the energy production (C category) (Fig. 2). Examples of essential genes common to these eight bacterial species encode ribosomal subunits, initiation factors of translation, cell division and chromosome replication, ATP synthase and respiratory chain subunits, peptidoglycan biosynthesis enzymes or enzymes involved in lipid A biosynthesis. Remarkably, among the latter are part of the genes of the *arnBCA_1_A_2_DarnT* cluster (BRPE64_RS06345-BRPE64_RS06375 in *C. insecticola*), encoding the biosynthesis of 4-amino-4-deoxy-L arabinose (Ara4N) and its transfer to the lipid A moiety of lipopolysaccharide (LPS) (Fig. 3B). This gene cluster was in an independent approach found to be essential for viability in *B. cenocepacia* strain K56-2 [41]. The Ara4N modification of LPS is a mechanism that mediates resistance against cationic antimicrobial peptides (AMPs) in many Gram-negative bacteria, including *Burkholderia* species [42]. In the γ-proteobacteria *Salmonella* and *Pseudomonas*, the Ara4N modification is not essential and not constitutively present in lipid A but is introduced upon sensing of AMPs in the environment [43–46]. In contrast, the essentiality of the Ara4N modification in *Burkholderia s.l.* spp. suggests that the Ara4N lipid A modification is constitutive. Indeed, the lipid A structure of *B. cenocepacia* and *C. insecticola* is modified with one or two Ara4N moieties [41, 46]. As demonstrated for *B. cenocepacia*, the Ara4N modification is required for LPS export by the Lpt transporter that transfers fully assembled LPS from the inner membrane to the outer membrane and that has an absolute specificity for Ara4N carrying LPS molecules [47].

### Conditional fitness defect genes

In a previous study, the comparison of the transcriptomes of the free-living and *R. pedestris* midgut-colonizing *C. insecticola* highlighted up or down regulated metabolic pathways [13]. Transporters or metabolic pathways of diverse sugars such as rhamnose and ribose, and sulphur compounds like sulphate and taurine (C_2_H_7_NO_3_S) were upregulated in the midgut-colonizing bacteria. Moreover, glycolytic pathways were down regulated and the gluconeogenesis pathway was upregulated. These data indicate that symbiotic bacteria could depend on particular sources provided by the insect. We used Tn-seq to identify key genes in *C. insecticola* for growth on these nutrients.

To mimic the specific nutrient conditions that can be found in the insect symbiotic organ, the *C. insecticola* transposon mutant library was grown in minimal media supplemented with different compounds. Glucose, 3-hydroxybutyric acid (HBA), mannitol, succinate, myo-inositol or rhamnose were used as the only carbon sources. Taurine was used as sole carbon source, or as sole nitrogen or sulphur source, or as sole carbon/nitrogen/sulphur source. Tn-seq data were obtained for each growth condition, and we used the resampling method analysis from the Transit software package to identify the fitness defect genes for each condition, using minimal medium with glucose or YG rich medium as the reference control condition (Supplementary Dataset s4). By these comparisons, we identified sets of condition-dependent fitness defect genes for each growth conditions (Fig. 4; Fig. s3; Supplementary Dataset s4). Examples are amino acid biosynthesis genes that are not fitness defect genes in rich medium, but are fitness defect genes in MM lacking any source of amino acids. Genes encoding glycolytic enzymes, like the phosphoenolpyruvate carboxylase *ppc* (BRPE64_RS03260), are fitness defect genes in MM supplemented with glycolytic substrates glucose, rhamnose, mannitol or myo-inositol but not with gluconeogenic substrates succinate, HBA and taurine. Inversely, the gluconeogenesis-specific genes *pps* (BRPE64_RS05810), encoding phosphoenolpyruvate synthase, and *fbp* (BRPE64_RS03750), encoding fructose-1,6-bisphosphatase, are fitness defect genes in MM with gluconeogenic substrates but not in MM with glycolytic substrates. *maeB* (BRPE64_RS11265), encoding the malic enzyme, is a fitness defect gene only in MM with succinate. Some genes appear as fitness defect in very specific conditions only. Examples are a contiguous cluster of nine genes encoding an ATP-binding cassette (ABC) transporter and involved in myo-inositol assimilation into the tricarboxylic acid cycle (TCA) cycle and gluconeogenesis/glycolysis (BRPE64_RS09045-BRPE64_RS09085) that has fitness defect only in the growth on myo-inositol condition. The gene BRPE64_RS07345, incompletely annotated as encoding a chloride channel, shows a fitness defect specifically in MM supplemented with taurine as carbon source, as nitrogen source, as sulphur source or as carbon/nitrogen/sulphur source. This suggests that the BRPE64_RS07345-encoded chloride channel protein is a taurine uptake transporter, which we named TauT. TauT is unlike known bacterial taurine transporters, which are ABC transporters [48] or Tripartite ATP-independent periplasmic (TRAP) transporters [49]. Interestingly, the mammalian taurine transporter is a chloride-dependent channel. Cluster BRPE64_RS16735 to BRPE64_RS16785 (11 genes) is specifically required for growth on rhamnose and codes for an ABC uptake transporter and the enzymes that incorporate rhamnose into pyruvate metabolism. The genes BRPE64_RS05370 and BRPE64_RS05375 encoding the subunits of 3-oxoacid CoA transferase allow to assimilate HBA into the TCA cycle and are specifically required for growth on HBA. Finally, genes BRPE64_RS02530 to BRPE64_RS02555 are specifically required for growth on mannitol and codes for an ABC uptake transporter.

**Figure 4.**
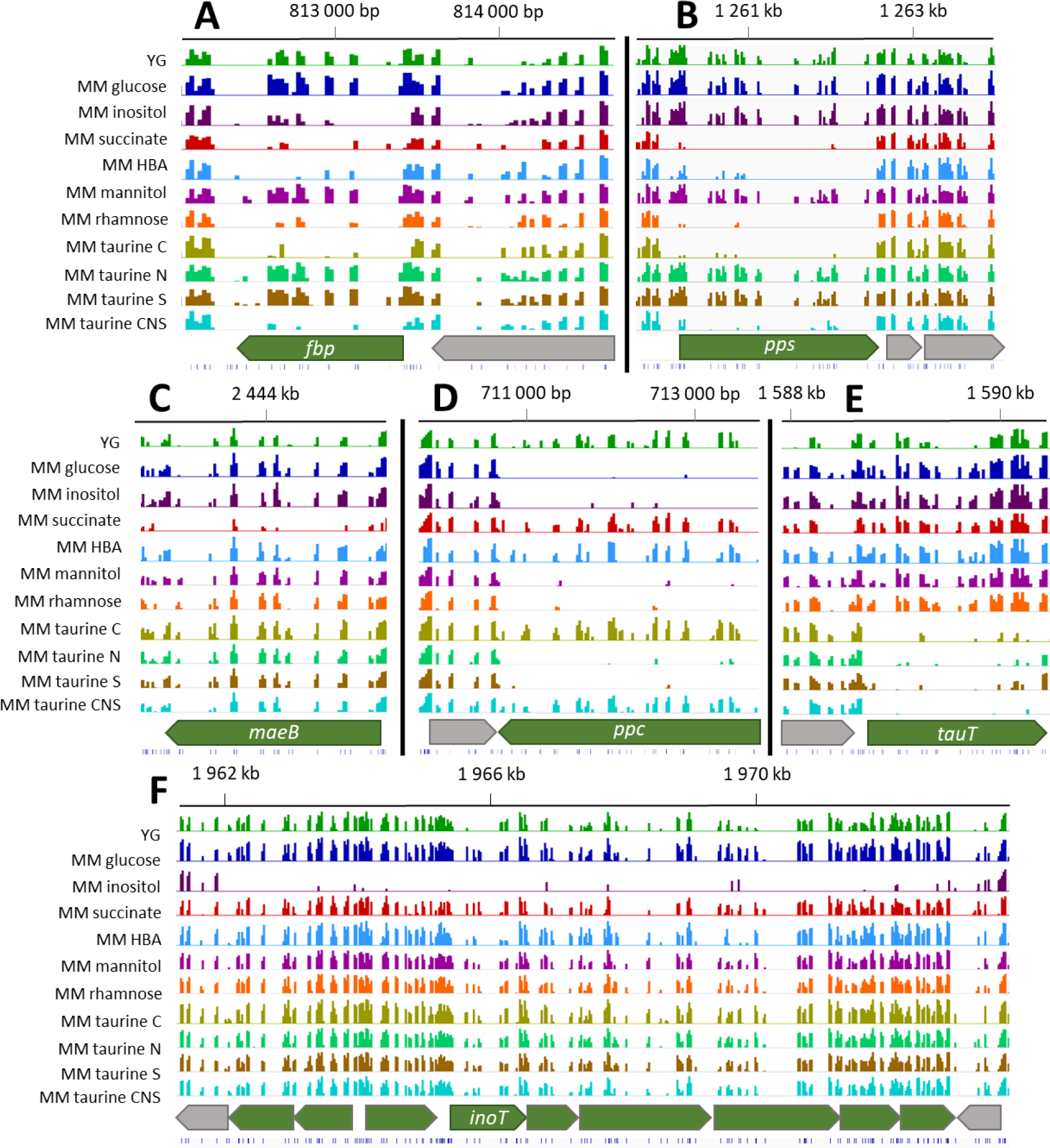
IGV plots of genomic regions carrying condition-specific fitness genes. **A.** Fructose-bisphosphatase encoding gene (*fbp*). **B.** Phosphoenolpyruvate synthase encoding gene (*pps*). **C.** Malic enzyme encoding gene (*maeB*). **D.** Phosphoenolpyruvate carboxylase encoding gene (*ppc*). **E.** Chloride channel protein and potential taurine transporter encoding gene (*tauT*). **F.** Myo-inositol utilisation genes including the ABC transporter permease (*inoT*). Tracks, from bottom to top: Position of TA sites (blue bars); Gene organization in the region of interest with fitness genes (in green) and their flanking neighbours (in grey); Histograms of insertion counts at TA sites for the indicated experimental conditions; Genome positions (in kb) on the chromosome 1. HBA, 3-hydroxybutyric acid; taurine C, taurine as carbon source; taurine N, taurine as nitrogen source; taurine S, taurine as sulphur source; taurine CNS, taurine as carbon, nitrogen and sulphur source.

### Confirmation of Tn-seq results by mutagenesis of selected conditional fitness-defect genes

To confirm the Tn-seq results, insertion mutants of *C. insecticola* were constructed in a set of six selected fitness genes (Fig. 4) for specific nutrient conditions, and their ability to grow on the corresponding nutrient was evaluated. These chosen genes were the above mentioned gluconeogenesis genes *fbp*, *pps* and *maeB*, the glycolysis-specific gene *ppc*, the newly discovered putative taurine transporter gene *tauT* and the myo-inositol transporter gene *inoT*.

Growth of the mutants in minimal medium with the relevant carbon sources and in the rich medium YG as a control allowed us to confirm the above Tn-seq results (Fig. 5). As expected and indicated by Tn-seq, all mutants grew well in YG. The growth of the *fbp* and *pps* mutants was affected in minimal medium with succinate, HBA, and taurine as carbon source; the *maeB* mutant was affected in minimal medium with succinate, the *ppc* mutant in minimal medium with glucose and myo-inositol, the *tauT* mutant in minimal medium with taurine as carbon source and the *inoT* mutant in minimal medium with myo-inositol.

**Figure 5.**
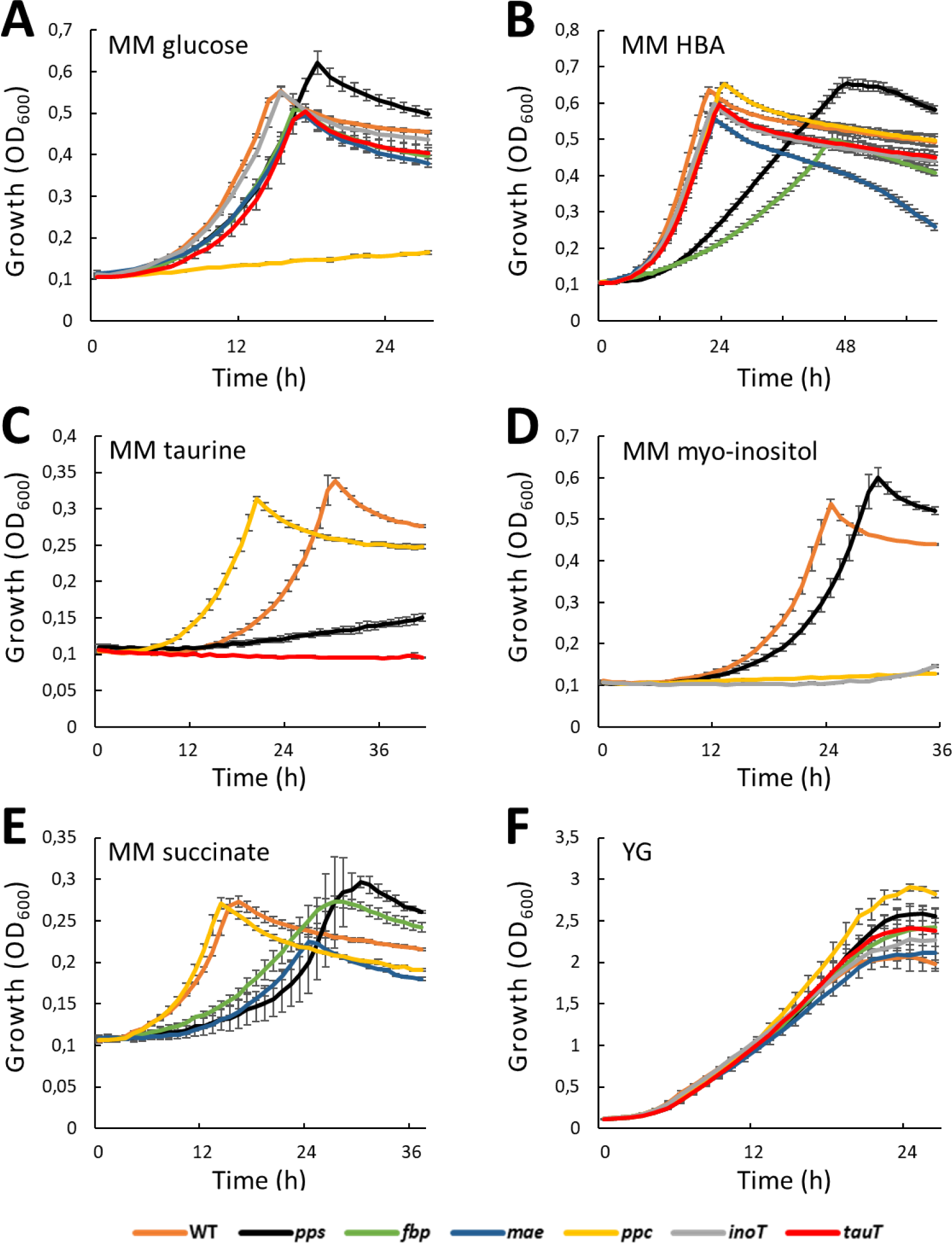
Growth curves of *Caballeronia insecticola* wild type (WT) and metabolic mutants in different media. **A.** Minimal medium with glucose. **B.** Minimal medium with 3-hydroxybutyric acid (HBA). **C.** Minimal medium with taurine as carbon source. **D.** Minimal medium with myo-inositol. **E.** Minimal medium with succinate. **F.** YG medium. X-axis, time of growth in hours; Y-axis, growth measured as OD_600_. Error bars are standard deviation.

Next, the capacity of the mutants to colonize the gut symbiotic organ in *R. pedestris* in mono-inoculation and in co-inoculation with the wild-type strain was tested (see Materials and Methods). The mutants were all able to colonize the insect symbiotic organ in mono-inoculation conditions (Fig. 6A). In co-inoculation, we calculated a competitive index to estimate the fitness difference between mutants and wild-type bacteria (Fig. 6B and C). For the three mutants with insertions in key enzymes of the gluconeogenesis, in particular the *fbp* and *pps* mutants, we observed that their capacity of colonization of the symbiotic organ was strongly affected in presence of the wild-type bacteria in co-inoculation experiments, implying that these three genes play a key role in the bacterial fitness for insect crypt colonization. The *tauT* and *inoT* mutants were also clearly outcompeted by the wild type although to a lesser extent than the three first mutants, suggesting that taurine and inositol are available nutrients in the crypt region. By contrast, the mutant in the glycolytic enzyme *ppc* was equally capable as the wild type to colonize the insect symbiotic organ, implying the absence of glycolytic carbon sources.

**Figure 6.**
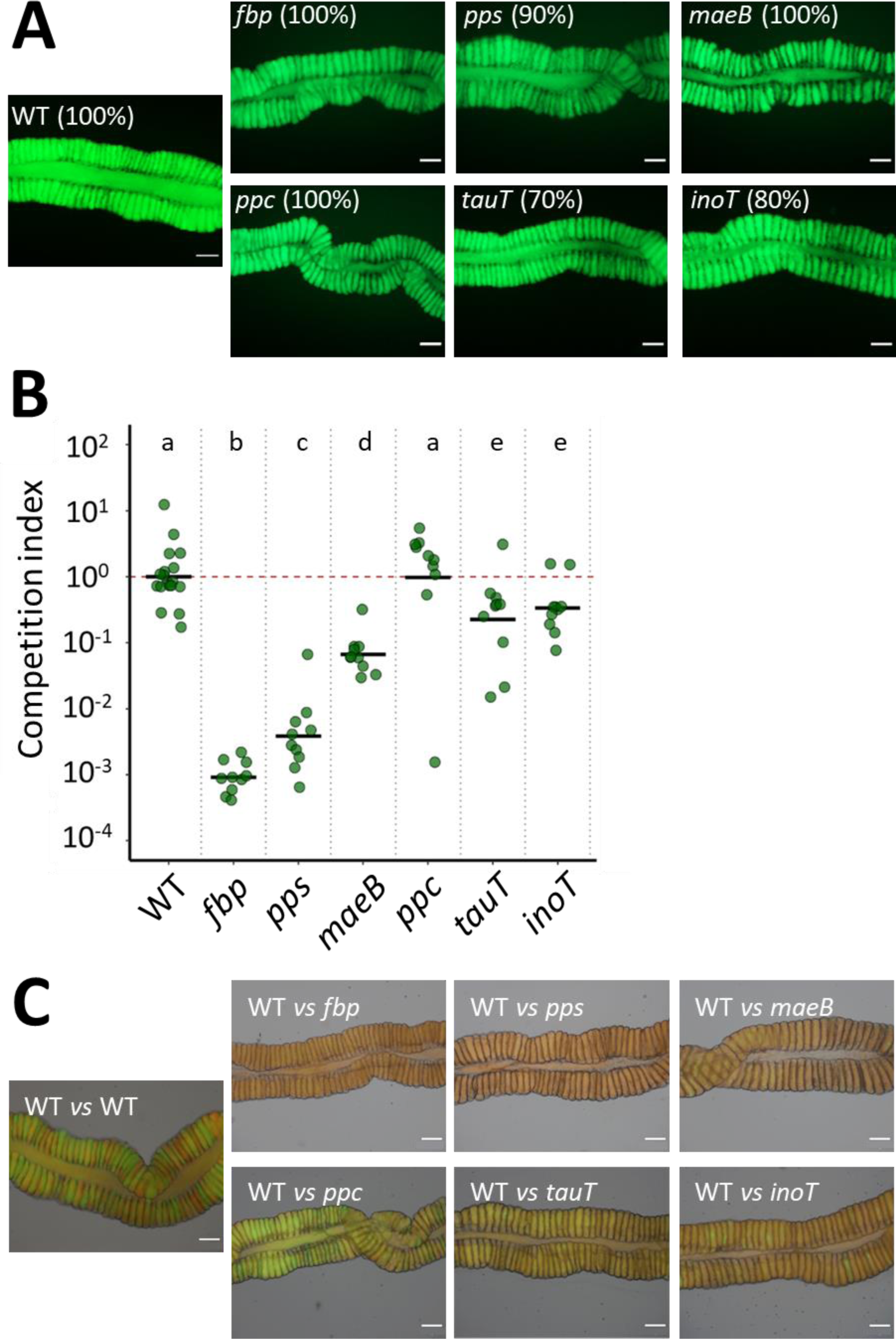
Ability of *Caballeronia insecticola* mutants to colonize the *Riptortus pedestris* M4 midgut region. Midguts were analyzed at 5 dpi. **A.** Colonization capacity of wild-type (WT) and mutant strains in single strain infection conditions were determined by microscopy observation with an epi-fluorescence microscope. One representative image is shown for each condition. Infection rate (%) indicates the proportion of infected animals with indicated strains (n=10). **B.** Colonization capacity of strains in co-infection conditions. *R. pedestris* was infected with an equal mix of mScarlett-labelled *C. insecticola* wild type and the indicated GFP-labelled wild-type or mutant strains. Relative abundance of the two strains in the M4 midgut regions at 5 dpi was determined by flow cytometry on dissected intestines. The competition index expresses for all samples the ratio of the indicated mutant to wild type, corrected by the ratio of the inoculum, which was in all cases close to 1 (mutant bacteria / wild-type bacteria) / (inoculum mutant bacteria / inoculum wild-type bacteria). Each dot represents the competition index in an individual and the mean per mutant is indicated by a horizontal black line (n=10). CI is competition index. Different letters indicate statistically significant differences (P<0.05). Statistical significance was analyzed by Kruskal–Wallis test, Dunn post hoc test and Benjamini-Hochberg correction. **C.** Microscopy observation with an epi-fluorescence microscope of competition assays between *C. insecticola* wild type (RFP) and mutants (GFP) as in panel B. One representative image is shown for each condition. Scale bars in panels A and C are 40 µm.

## Discussion

Discovering the essential genome of bacteria is an important step towards the understanding of the principal biological functions determining their life styles (3,4]. The availability of a transposon mutant library of *C. insecticola* and the genome-wide identification of essential and conditionally fitness genes constitutes therefore a step towards the characterization of the fitness landscape of this bacterium in its different living environments, which include persistence in soil, colonization of the symbiotic organ of insects as well as interactions with other organisms present in the natural environment of this bacterium like soil microbes and plants.

Chromosome 1 carries the large majority of genes conserved between *Burkholderia s.l.* species, while the other replicons are more enriched in genes with limited species distribution [50]. Accordingly, we found that nearly all the essential genes of *C. insecticola* are located on chromosome 1. This observation is moreover in agreement with the plasmid origin of the other two chromosomes, which therefore can be considered as chromids. Probably, ancestral plasmids acquired genes, including essential ones, from chromosome 1 after their capture by the ancestor of *C. insecticola*, giving rise to the present essential chromids. Since chromosome 3 contains only two essential genes besides the replication/partitioning genes, suggesting that, after translocation of these two essential genes to the chromosome 1 or 2, this replicon could be entirely cured from the *C. insecticola* genome. In agreement, it was shown that the chromosome 3 in *Burkholderia cenocepacia* strains can be efficiently cured [51, 52]. The absence of essential genes in the plasmids is in line with the demonstration that plasmid 2 can be removed from the bacterium without affecting its fitness [13] and further suggests that the large plasmid 1 can be removed as well without affecting viability. A reduced-genome engineered strain lacking chromosome 3, plasmid 1 and plasmid 2 could be a powerful platform that can be used for gene discovery.

Among the 498 essential genes of *C. insecticola*, we found only a surprisingly small portion of them conserved among related species from the *Burkholderia s.l.*. However, despite the fact that essential genes code for fundamental cellular functions, essential gene sets are known to be specific to each bacterium, and they can even vary among strains belonging to the same bacterial species, similarly as we find here for the three analyzed *B. cenocepacia* strains [53–55]. Non-orthologous gene displacement is one explanation that has been put forward for the absence of conservation of essential genes among bacteria. This concept proposes that essential pathways or genes are replaced or co-exist with functional equivalents with no DNA homology and different evolutionary origin [55]. Gene duplication on the other hand could render genes non-essential because of redundancy while the encoded function remains essential. Gene duplication is observed only for a few genes that are non-essential in *C. insecticola* but essential in related species, suggesting that non-orthologous gene displacement is accounting for a large part of the differences among the essential gene sets in *Burkholderia s.l.* species. Moreover, it should be mentioned that different Tn-seq techniques and bioanalysis tools were employed for the determination of the essential gene sets of these eight species. This heterogeneity in analysis methods may create biases in the identification of common essential genes and potentially their number is in reality higher than that we determined here.

We showed here that the transposon library in *C. insecticola* in combination with the Tn-seq method provides robust fitness data at the whole genome level and that it can be used efficiently to identify conditionally essential genes in this species. We identified genes that are specifically essential for growth in media with gluconeogenic or glycolytic carbon sources, with rhamnose, mannitol, myo-inositol and taurine and confirmed their metabolic function by mutagenesis for a subset of them. Our Tn-seq study thus contributed to annotate genes with a previously unknown role as exemplified by the chloride channel required for taurine utilisation.

The phenotypic characterization of constructed mutants further revealed that the *C. insecticola* bacteria in the gut are fed by the insect with gluconeogenic carbon sources, including taurine, as well as with myo-inositol but not with glycolytic nutrients. It appeared that if the use of gluconeogenic substrates is crucial for an efficient colonization, the capacity to utilize individual compounds like taurine or myo-inositol has a less strong impact because mutants, which are unable to import these molecules, are still able to partially colonize the insect gut in competition with the wild type. It should be noted that phosphoenolpyruvate carboxylase encoded by *ppc* is required for growth on myo-inositol as a sole carbon source (Fig. 6D) while *ppc* is not required for gut colonization. Therefore, myo-inositol in the gut is unlikely to fuel the TCA cycle via *ppc* but is rather assimilated into a different pathway. Moreover, all the tested mutants were able to colonize the crypt region in the absence of competition with wild-type bacteria. Together, this suggests that the insect is providing multiple but specific nutrients to the bacteria. Thus, the nutritional exchange in the symbiosis between *C. insecticola* and *R. pedestris* is complex [13]. Our study demonstrates that Tn-seq can contribute to dissect this process in detail. An exciting possibility, worth to be investigated in the future, is that the feeding of the crypt bacteria with specific nutrients tailors the symbiont metabolic fluxes towards an optimized production of particular beneficial metabolites for the host.

In conclusion, the Tn-seq approach is a powerful tool for whole genome genetic screens in *C. insecticola* and future Tn-seq experiments in other *in vitro* conditions or in natural environments, like in the insect gut, soil, rhizosphere or co-cultivation with other microorganisms would strongly help to better understand the different lifestyles of this bacterium. Defining the genetic repertoires that determine the fitness of *C. insecticola* in these environments will highlight pre- and post-adapted traits of the *Caballeronia* symbiont to the insect’s gut environment.

## Acknowledgements

This work beneficiated from financial support by Saclay Plant Sciences-SPS, by the Agence Nationale de la Recherche, grant ANR-19-CE20-0007, and by a JSPS-CNRS Bilateral Open Partnership Joint Research Project and a CNRS International Research Project to Y.K. and P.M. Y.K. was supported by the Ministry of Education, Culture, Sports, Science, and Technology (MEXT) KAKENHI (18KK0211, 21K18241, 22H05068). T.O. was supported by the JSPS Research Fellowship for Young Scientists (14J03996, 20170267 and 19J01106). J.L., G.L. and R.J. were supported by Ph.D. fellowships from the French Ministry of Higher Education, Research, and Innovation. Tn-seq sequencing and data treatment were performed by the I2BC high-throughput sequencing facility, supported by France Génomique (funded by the French National Program Investissement d’Avenir ANR-10-INBS-09).

## Competing Interests

The authors declare no competing interests.

## Data Availability Statement

Tn-seq sequencing data were deposited in the SRA (BioProject accession n° PRJNA1004562 and PRJNA890438). All other data generated or analyzed during this study are included in this published article and its supplementary information files.

## Author contributions

P.M. and T.O. designed the study, planned the experiments and supervised the project. J.L., R.J., D.N. performed Tn-seq. G.L., R.C. and T.T. made mutants. R.J. performed *in vitro* assays. G.L., A.Y., R.C and T.O. performed insect experiments. R.J, G.L., T.O., Y.K, and P.M. analyzed data. R.J. and P.M. wrote the manuscript with input from T.O. and Y.K. All authors provided critical feedback and helped to shape the manuscript.

## Supplementary information

### Content

Supplementary Figures s1 to s3

Supplementary Table and Datasets 1 to 4

Supplementary Text

**Supplementary Figure s1.**
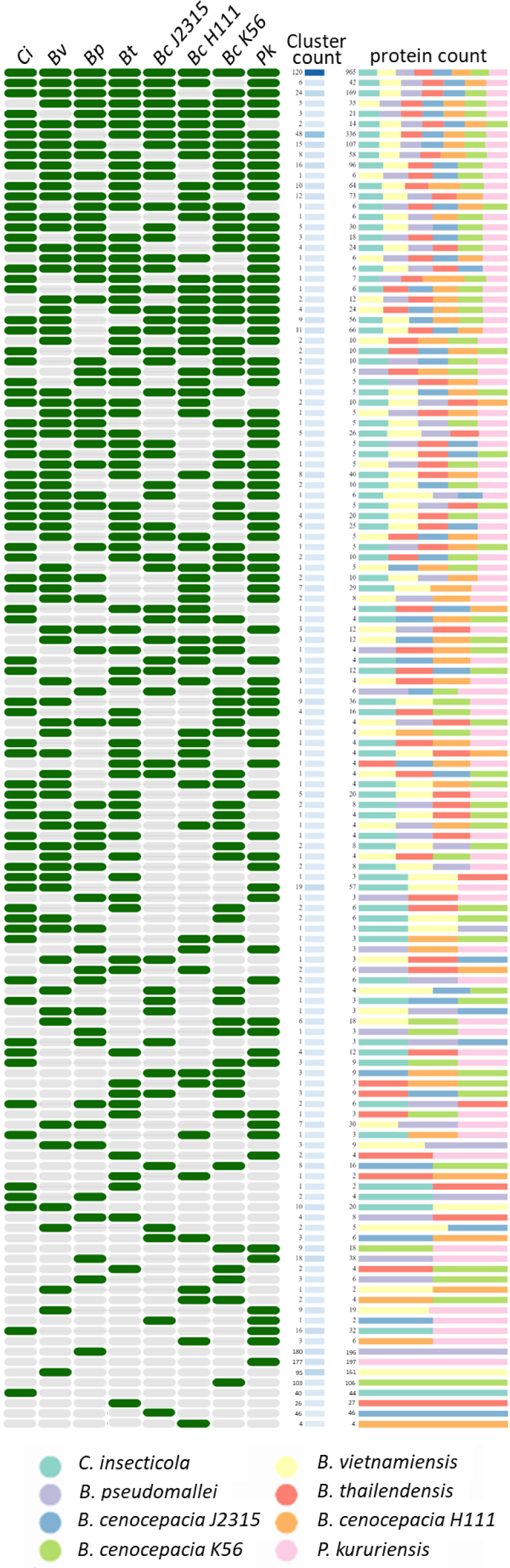
Homology-based comparison of essential genes shared between eight *Burkholderia s.l.* species using OrthoVenn2. The essential gene sets of the eight indicated species are pooled and orthologous groups (clusters) are created. The presence of clusters in the species are indicated by green cells and absence with grey cells. Between species shared cluster numbers are indicated in the cluster count column and by the colour code from dark to light blue of the corresponding cell. The protein count indicates per cluster the number of proteins present in each species according to the colour key.

**Supplementary Figure s2.**
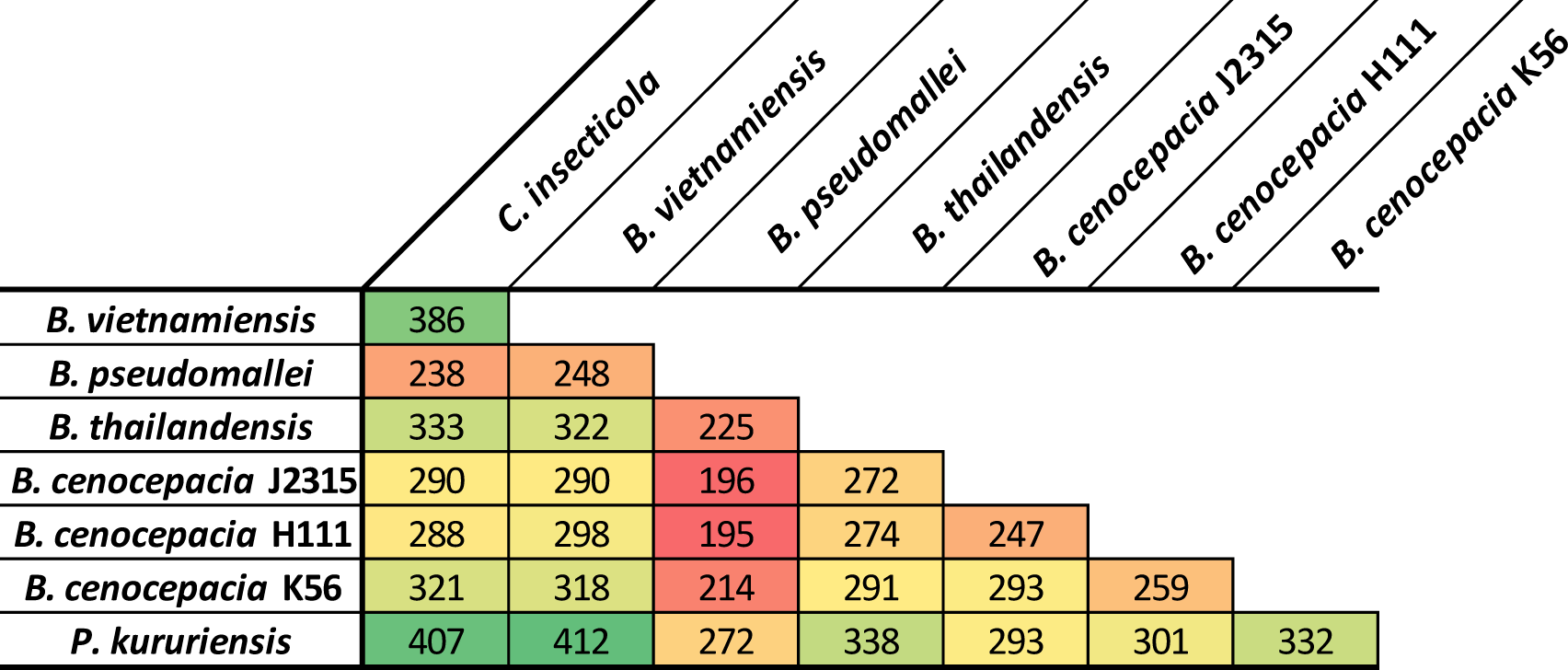
Pairwise comparison of essential gene sets in eight analysed *Burkholderia s.l.* species.

**Supplementary Figure s3.**
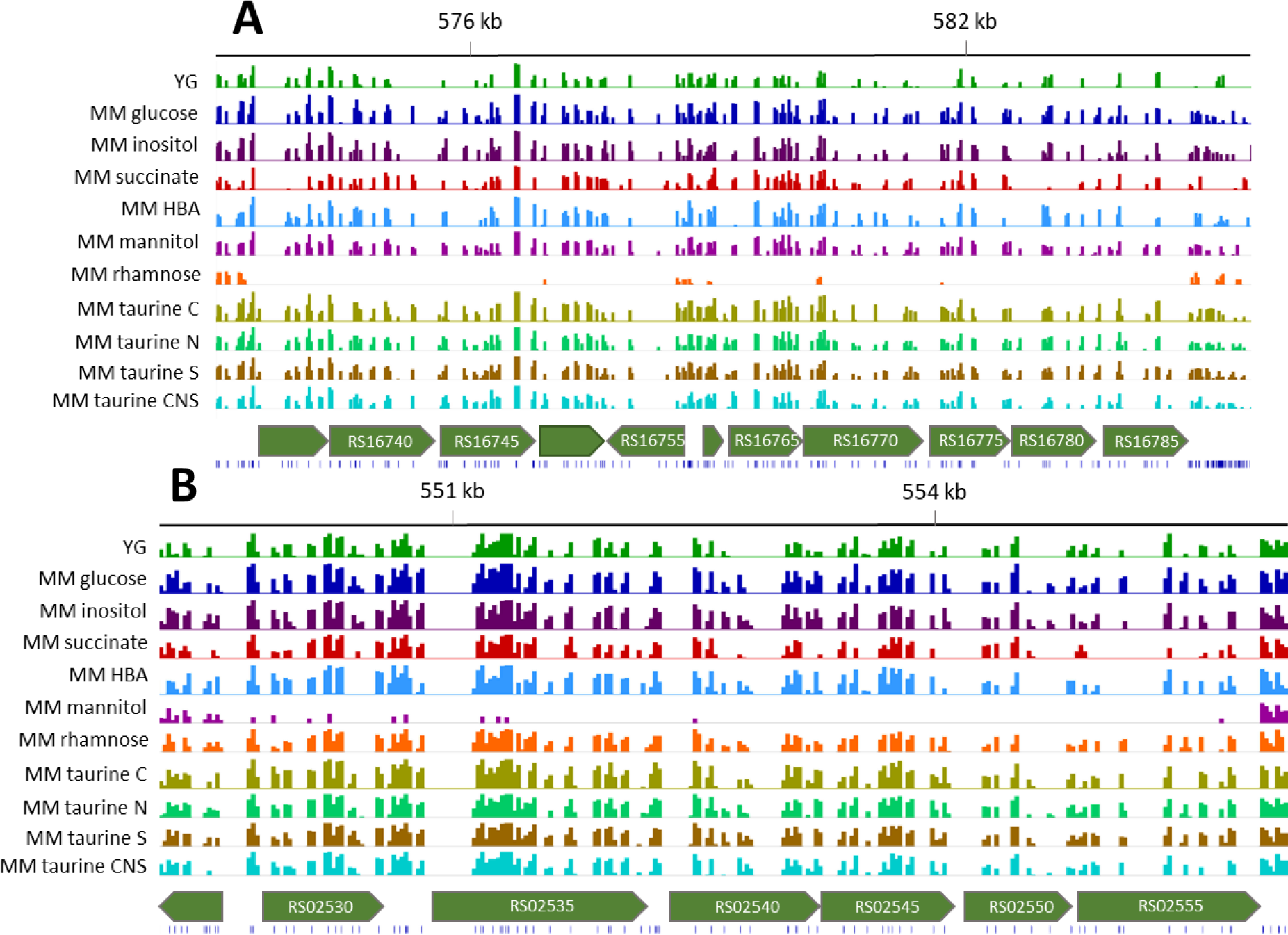
IGV plots of genomic regions carrying condition-specific fitness genes. **A.** Rhamonose utilisation genes. **B.** Mannitol utilization genes. Tracks, from bottom to top: Position of TA sites (blue bars); Gene organization in the region of interest with fitness genes (in green) and their flanking neighbours (in grey); Histograms of insertion counts at TA sites for the indicated experimental conditions; Genome positions (in kb) on the chromosome 1. HBA, 3-hydroxybutyric acid; taurine C, taurine as carbon source; taurine N, taurine as nitrogen source; taurine S, taurine as sulphur source; taurine CNS, taurine as carbon, nitrogen and sulphur source.

## Supplementary Table and Datasets

**Supplementary Table s1.** Primers used in this study.

**Supplementary Dataset s1.** Composition of minimal media used for Tn-seq screens and growth curves.

**Supplementary Dataset s2.** The essential genes of *C. insecticola* identified by Transit analysis and results of HMM analyses for the YG, MM glucose and MM succinate conditions.

**Supplementary Dataset s3.** Essential genes in *C. insecticola* common between the eight *Burkholderia s.l.* species and Orthovenn2 clusters.

**Supplementary Dataset s4.** Transit resampling analysis of Tn-seq data from growth on various nutrients.

## Supplementary text

### Generation of a *C. insecticola* RPE75 *Himar1* transposon library

The *E. coli* MFDpir strain carrying the plasmid pSAM_EC with a modified *Himar1* mariner transposon, containing the *nptII* kanamycin resistance gene [1] was used as a donor strain for transposon mutagenesis. The donor strain *E. coli* MFDpir pSAM_Ec and the recipient strain *C. insecticola* RPE75 were grown in 50 mL liquid cultures until exponential growth phase with a final OD_600nm_≈1. Bacteria were washed twice by centrifugation of cultures at 4000 rpm for 10 minutes at 4°C and resuspension of pellets in fresh medium. Final pellets were resuspended in 1 mL fresh medium to obtain an OD_600nm_≈50. For conjugation, the donor and the recipient strain were mixed at a 1:1 ratio, spotted per 100 µL on YG agar plates supplemented with 300 µg/mL DAP and plates were incubated at 28°C. After 1 hour of incubation, 1 ml of YG medium was added per conjugation spot to recover bacteria. Dilution series of this bacterial mix were plated on a selective medium carrying Rif (selection of RPE75) and Km (selection of the transposon) and subjected to colony forming units (cfu) counting to assess the number of independent bacterial mutants obtained by the mutagenesis, which was estimated to be about 2.5×10^8^ clones. In parallel, the totality of the remaining bacterial suspension was spread on 100 YG agar plates (100 µL per plate) supplemented with Rif and Km to obtain the *C. insecticola* transposon mutant population. After 2 days of incubation at 28°C, the transposon library was resuspended from the agar plates in fresh liquid YG medium. The suspension was adjusted to 20 % glycerol, aliquoted per 1 mL and stored at −80°C. The titer of this library was about 2×10^10^ cfu/mL for a total of 127,5 mL.

Before further use, a quality control was performed on the library. The presence of the mariner transposon and the absence of the transposon donor plasmid pSAM_Ec were verified by PCR on 20 randomly selected clones of the obtained transposon library. The transposon borders of these clones were amplified by PCR as described below and inserted into the pGEM-T Easy plasmid (Promega). For each of these 20 pGEM-T constructs, 10 clones were sequenced. This verification confirmed that each initial *C. insecticola* clone carried a single *Himar1* transposon insertion, that the two borders of the transposon were obtained and that the each of the 20 randomly selected clones carried a transposon in a distinct genomic location, distributed over the genome.

### DNA extraction and preparation of the high-throughput sequencing libraries

Genomic DNA was extracted from the bacterial pellets using the MasterPure™ Complete DNA and RNA purification kit (Epicentre) according to the manufacture’s instructions. Samples were cleaned-up by removing RNA according to instructions. Samples of 10 µg DNA were digested for one hour at 37°C with 1 µL of *Mme*I enzyme (2000 U/mL, New England BioLabs), in a total volume of 250 µL mix supplemented with 25 µL of 10X CutSmart buffer (New England BioLabs) and 10 µL of S-adenosine-methionine (1.5 mM, New England BioLabs). Subsequently, 1 µL of FastAP Thermosensitive Alkaline Phosphatase (1 U/µL, ThermoScientific) was added to the digestion mixes and samples were incubated for one additional hour at 37°C. The enzymes were then heat-inactivated at 75°C for 5 minutes. Digested DNA samples were purified from the reaction mix using the QIAquick PCR purification kit (QIAGEN). 700 ng of digested DNA was ligated to specific barcoded adaptors (5 µM) (Table s1) using T4 DNA ligase (1 U/µL, ThermoScientific) in a final volume of 20 µL and incubated overnight at 16°C. The double stranded adaptors were prepared beforehand by mixing 25 µL of each corresponding single stranded primer at 200 µM (Table 1) and 1 µL of TrisHCl (100 µM, pH 8.3), denaturing the primers in the mixture at 92°C for 1 min and promoting the annealing of the complementary primers by gradual cooling of the samples (2°C per min) in a PCR thermocycler. Transposon borders were subsequently amplified by PCR from the adapter-ligated DNA samples using 1 µL of them as template. The PCR was performed for 22 cycles using the EuroBio Taq polymerase (5 U/µL, reference GAETAQ00-4W) in a final volume of 20 µL according to the manufacturer’s instructions, with 0.5 µM of the forward P7 Illumina primer and 0.5 µM of the reverse P5 Illumina primer (Table s1 for primer sequences). The amplified products (130 bp) were separated on a 2.5 % agarose gel by electrophoreses and purified from the gel using the QIAquick gel extraction kit (QIAGEN). The concentration and the quality control of these Tn-seq Illumina sequencing library samples were assessed using Qubit fluorometric quantification (ThermoFisher) and a Bioanalyzer instrument (Agilent), respectively.

### Sequencing and sequence data treatment

Up to 20 Tn-seq samples were mixed in equimolar amounts and sequenced by an Illumina NextSeq 500 instrument with 2 x 75 paired-end run at the I2BC sequencing platform (CNRS Gif-surYvette, France). The generated data were demultiplexed using bcl2fastq2 software (bcl2fastq v2.15.0; Illumina, San Diego, USA) and FASTX-Toolkit (http://hannonlab.cshl.edu/fastx_toolkit). The 3’ transposon sequence was trimmed using Trimmomatic [2], and reads with a length of 75 nucleotides were removed (reads without the transposon insertion). After the trimming step, reads with a length between 19 and 23 bp were reverse-complemented and only the reads starting with TA dinucleotides were mapped using Bowtie (bowtie-1.1.2) [3,4] to the reference genome of *C. insecticola* [5] (accession n° NC_021287.1 (chromosome 1), NC_021294.1 (chromosome 2), NC_021288.1 (chromosome 3), NC_021289.1 (plasmid 1), NC_021295.1 (plasmid 2)). BAM output files were sorted with Samtools (http://www.htslib.org/). FeatureCounts [6] was used to evaluate the number of reads per gene. BAM output files were converted with Samtools on the Galaxy server (https://usegalaxy.org/) into non-binary SAM files, the appropriate format to use for further analysis.

### Identification of (conditionally) essential genes by Transit software

Tn-seq sequencing data was handled by TRANSIT Version 3.2.0. It provides an easy to use graphical interface and access to several different Tn-seq data analysis methods that allow the user to determine essentiality within a single condition (Hidden Markov Model analysis) as well as between two conditions (Resampling analysis) [7].

The Hidden Markov Model (HMM) analysis from Transit software allows determining fitness of each TA site and each gene of the genome of interest in a single condition. By this analysis, genes are classified as non-essential (NE), growth defect (GD), growth advantage (GA) or essential (ES). The selected HMM parameters in TRANSIT were 10 % cutoff, TTR normalization and sum of replicates. The resampling analysis from Transit software compares the number of transposon insertions in a control condition to the number of transposon insertions in the experimental condition for each gene of the genome of interest. For each gene, a *p*-value to assess the significance of the difference with the control condition and a log2-fold-change expressing the importance of these differences is obtained. The parameters used in the resampling analysis were 10 % cutoff, 10 000 samples, pseudo-count of 5 and TTR normalization. Selected cut-off values for significant fitness change were |log2-fold-change|>1.58 and *p*<0.05. Each experimental condition was compared to the MM with glucose as the reference condition.

The Integrative Genomics Viewer (IGV) software, an interactive tool for the visual exploration of genomic data, was used to visualize the number of insertions per insertion sites in specific genes of interest [8].

### Creation of fluorescent protein tagged *C. insecticola* strains

A mScarlett-I-tagged strain of *C. insecticola* was created by introducing a Tn7-Scarlet transposon. The Tn7-Scarlet donor strain S17-1λpir.pMRE-Tn7-135, the helper strain WM3064.pUX-BF13 and *C. insecticola* were grown overnight in LB with DAP if appropriate and YG respectively. Overnight cultures were diluted in 10 mL fresh medium using 0.2 mL of the overnight cultures and grown at 180 rpm until reaching a final OD_600nm_<1. The cultures were washed twice in fresh YG medium without antibiotics by centrifugation at 4000 rpm for 10 minutes. Bacterial pellets were resuspended in fresh YG medium to obtain a final OD_600_≈5 to 10. Then the donor, helper and *C. insecticola* strains were mixed at a 1:1:1 ratio. 150 µL of the conjugation mix was spotted on YG agar plates with DAP and 0.1 % of L-arabinose to induce transposition. After overnight incubation at 28°C the spots on the plates were resuspended in 1 mL of YG medium. Ten-fold dilution series were made and 50 µL of each dilution were plated out on YG plates complemented with Cm. DAP was not added to counterselect the *E. coli* strains from the mixture. Selected colonies were screened for mScarlet-I expression by UV light illumination and they were purified again on YG complemented with Rif and Cm.

### Insect rearing and inoculation tests

The bean bug, *R. pedestris,* was originally collected from Japan, from soybean field in Tsukuba, in 2007. The insects are maintained in the laboratory and are reared in plastic boxes at 25°C under a long-day regimen (16 h light, 8 h dark) and fed with soybean seeds and distilled water containing 0.05 % ascorbic acid (DWA). After hatching, insect eggs were transferred into sterile Petri dishes. After two days, at the second larval stage, water was removed to make insects thirsty, which facilitates the subsequent ingestion of administered bacteria. After overnight starvation, a bacterial suspension of the tested *C. insecticola* strain adjusted at 10^7^ cfu/mL in sterile distilled water was provided for infection of the second instar nymphs. For co-inoculation experiments, two bacterial strains were mixed together at a 1:1, each adjusted before to 10^7^ cfu/mL.

At three and five days post inoculation, insects, at the stage of the end of the second instar nymphs or the third instar, respectively, were dissected. Dissections were performed in sterile PBS (137 mM NaCl, 8.1 mM Na_2_HPO_4_, 2.7 mM KCl, and 1.5 mM KH_2_PO_4_, pH 7.5) containing 0.01 % of Tween 20 under a Stemi 508 binocular microscope equipped with an Axiocam 208 color camera (Zeiss). The M4 region of the midgut was collected using fine forceps and assembled on glass slides for microscopy observations (Nikon Eclipse 80i). The colonization rate of the inoculated insects was estimated by fluorescent signal detection of colonizing bacteria with GFP or mScarlett-labelled fluorescent proteins. Merged fluorescence pictures were obtained with GIMP version 2.10.32. Then M4 region samples were homogenized in PBS solution and bacteria in suspension were counted by flow cytometry, using the fluorescent tags to determine the relative abundance of the two inoculated strains. Flow cytometry was performed on a CytoFlex S instrument operated by CytExpert 2.4.0.28 software (Beckman Coulter). Gating by the forward-scatter (FSC) and side scatter (SSC) dot plot permitted to collect signals specifically derived from bacteria. Doublets were discarded using the SSC_Area-SSC_Height dot plot. GFP fluorescence was excited by a 488-nm laser and collected through a 525/40 nm band pass filter; RFP fluorescence was excited by a 561-nm laser and collected through a 610/20 nm band pass filter. Data acquisition for a total of 50 000-100 000 bacteria was performed for each sample. Thresholds for considering positive events for GFP and RFP were determined using non-fluorescent control bacteria. Data was treated by CytExpert and Excel software. Statistical analysis were performed in R using a Kruskal wallis test, with pairwise Wilcoxon test as post hoc test and the Benjamini-Hochberg p.adjust method (*p* <0.05).

## Notes

### Competing Interest Statement

The authors have declared no competing interest.

